# Structural basis of hypoxic gene regulation by the Rv0081 transcription factor of *Mycobacterium tuberculosis*

**DOI:** 10.1101/465575

**Authors:** Ashwani Kumar, Swastik Phulera, Arshad Rizvi, Parshuram Sonawane, Hemendra Singh Panwar, Sharmistha Banerjee, Arvind Sahu, Shekhar C. Mande

**Author notes:** To whom correspondence should be: Shekhar C. Mande, National Centre for Cell Science, NCCS Complex, SP Pune University Campus, Ganeshkhind, Pune - 411 007, INDIA.

## Abstract

The transcription factor Rv0081 of *M. tuberculosis* controls the hypoxic gene expression and acts as a regulatory hub in the latent phase of tuberculosis infection. We report here the crystal structure of Rv0081 at 3.3 Å resolution revealing that it belongs to the well-known ArsR/SmtB family proteins. ArsR/SmtB family transcriptional repressors exert gene regulation by reversible metal binding. Hypoxia in general is sensed by bacterial transcriptional regulators via metals or Cys-mediated thiol switches. Oxygen sensing typically leads to transcriptional repressor changing its conformational state with altered DNA-binding property under different oxygen levels. Surprisingly Rv0081 neither has a metal binding domain nor does it possess Cys residues suggesting an alternate mechanism of gene regulation. Our structural analysis identified Ser 48, Ser 49, Ser 52 and Gln 53 as potential residues of Rv0081 involved in DNA binding. We probed DNA-binding of Rv0081 with electrophoretic mobility shift assay (EMSA) as well as surface plasmon resonance (SPR), where the Alanine mutants of these residues showed diminished DNA binding. Similarly, Aspartate mutants of these Ser residues was shown to fail to bind to DNA. Since, phosphorylation of various regulatory proteins is one of the important controlling mechanisms, we expected the role of Ser-phosphorylation of Rv0081 in hypoxic condition. Probing Rv0081 with anti-phosphoserine antibodies in *M. tuberculosis* cell lysate showed marked enhancement in the phosphorylation of Rv0081 protein under hypoxia. Overall, our structural and biochemical analysis provides the molecular basis for the regulation of Rv0081 in the latent phase of tuberculosis infection.

**IMPORTANCE:** Tuberculosis is one of the deadliest infectious diseases caused by the bacterium *Mycobacterium tuberculosis*. In about 90% of the infected people, *M. tuberculosis* exists in a dormant or a latent stage which can be reactivated in favorable conditions. Hypoxia (low oxygen pressure) is one of causes of dormancy. Understanding hypoxic gene regulation in *M. tuberculosis* is therefore an important step to understand latency. Rv0081 is a transcriptional regulator of genes expressed during hypoxia. In order to understand the mechanism by which Rv00081 regulates gene expression during hypoxia, we have solved the crystal structure of Rv0081 and identified amino acid residues which are critical in its transcriptional regulator activity. The crystal structure is suggestive of mechanism of gene regulation under hypoxia.

## INTRODUCTION

A significant aspect of the pathogenesis of *Mycobacterium tuberculosis* (Mtb), the tuberculosis (TB) causing bacterium, is the latency stage, where the bacteria persist in an infected person asymptomatically. Latent TB is characterized by the presence of non-replicating persistent bacteria that successfully evade host immunity and survive inside the hostile macrophage environment (1, 2). Apart from slow replication, these mycobacteria are reported to develop resistance to anti-mycobacterial agents. While almost all the available drugs act on the actively dividing bacteria, the prolonged treatment of 4-6 months, if not more, is therefore followed to eliminate the non-replicating subpopulations that continue to thrive and lead to reduced susceptibility to standard TB drugs. Reactivation of the latent bacteria contributes to an active infection and consequently spread in human population, making TB control and eradication highly challenging. Understanding the processes of Mtb latency and reactivation is therefore crucial for controlling the spread of tuberculosis (3). Latency and reactivation of Mtb has been experimentally demonstrated to be intimately related to oxygen tension. A variety of experiments involving *in vitro* cell culture to animal models, supported by clinical observations in humans, have led to establishment of models that mimic clinical Mtb latency (3–6).

It has been observed that progressively increasing hypoxic condition, where mycobacteria are grown as sealed, standing cultures (the Wayne model), induced similar physiological, transcriptional and proteome changes as exhibited by latent mycobacteria (7, 8). Subsequently, re-aeration, in which mycobacteria grown under hypoxic conditions are transferred to aerated, shaking cultures, could mimic reactivation *in vitro* (6, 9, 10). Such cell culture based models have been used to understand physiology of Mtb under hypoxic conditions. Hypoxia sensing mechanism of Mtb has been delineated through genetic manipulations and comparative transcriptomics (10–13).

Hypoxia activated DosR/DosS (DevR/DevS) and MprAB two-component signal transduction regulatory system were found to regulate nearly 48 genes implicated in dormancy and granuloma like conditions (14–16). Later, the DosR-dependent regulon was observed to be induced in response to nitric oxide, carbon monoxide, low pH, in macrophages and in both early and late mouse infections, establishing beyond doubt the significance of this regulatory operon in dormancy (18). Subsequently, it was shown that besides DosR-mediated initial hypoxic response, sustained hypoxia spanning to 4-7 days induced a set of 230 genes, referred to as Enduring Hypoxic Response (EHR), that facilitated the shift to a persistent, metabolically inactive, but viable state (6, 19). A high throughput study by Galagan *et al*. combining ChIP-Seq data with system-wide transcriptomics, proteomics, metabolomics and lipidomics during hypoxia and re-aeration identified several transcriptional regulators outside that of the DosR or EHR response (20). This study identified Rv0081, a part of the DosR regulon, as a major regulatory hub for multiple hypoxia induced pathways. This supported our earlier simulation studies where we had predicted DosR and DosS, along with Rv0081 as important regulators of latency (21). Thus, Rv0081 is likely to be a major factor mediating transcriptional changes during transition between normoxic and hypoxic conditions.

Rv0081, the first gene of the operon *rv0081-rv0088*, belongs to the ArsR/SmtB family of prokaryotic metalloregulatory transcriptional repressors. This family of repressors is known to regulate the expression of genes linked to heavy metal ion stress including those involved in metal uptake, efflux or detoxification (22). Rv0081 was established experimentally as a repressor where Rv0081 self regulates its expression by binding to an inverted repeat element upstream to the locus *rv0081-rv0088* (23). Besides self-regulation, Rv0081 was shown to co-regulate the locus *rv0081-rv0088*, the genes of which are predicted to encode formate hydrogenlyase (FHL) enzyme complex. FHL is involved in degradation of formic acid, which is formed fermentatively under anaerobic conditions, into carbon dioxide and hydrogen (24, 25).

Formate is one of the common fermentation products in many organisms in their anaerobic phase, generated when the glycolytic product pyruvate is cleaved to acetyl-CoA and formate by pyruvate-formate-lyase (26). Secretion of formate in these organisms is essential for redox homeostasis. Apart from redox balance formate also serves as a source of energy for many microorganisms during aerobic or anaerobic respiration (27). The oxidation of formate produces a pair of electrons, a proton, and a molecule of carbon dioxide by formate dehydrogenase (FDH) complex which is located in the cytoplasmic membrane. This reaction typically occurs in the periplasm to avoid acidification of the cytoplasm, and electrons are transferred through the FDH complex to menaquinone along with protons from the cytoplasmic side of the inner membrane (27). Further, due to its low redox potential formate can also be oxidized by other enzyme systems. Notable examples of such systems are anaerobic reduction of fumarate and nitrate/nitrite. The cells thus have a myriad of choices with regards to formate and depending on the prevailing physiological conditions, choose to secrete it, oxidize it aerobically, use it as a reductant in anaerobic respiration with a variety of terminal reductases, or convert it to CO_2_ and H_2_ by formate hydrogen lyase complex (FHL). Another important aspect related to formate is its ability to serve as the hydrogen donor for ribonucleoside reductases (RNR). RNRs are responsible for the production of deoxyribonucleotides and typically accept electrons from either the thioredoxin or the glutaredoxins (28). In the case of Mtb, RNR accepts electrons from a thioredoxin like protein called NrdH (29). Formate related metabolic regulatory network called as formate regulon thus exists, with formate as the central player.

As mycobacteria would require shifting metabolism to anaerobic conditions inside granuloma during latency, Rv0081 might act as one of the key regulators deciding the fate of mycobacterial latency. By virtue of controlling the expression of genes on the *rv0081-rv0088* operon, Rv0081 might be required to sense oxygen levels in the cells, and thereby trigger gene expression under low oxygen levels. Wide ranging oxygen sensing mechanisms in prokaryotic cells have been described earlier, which include those mediated by metals (30), iron-sulfur clusters (31) or by means of Cysteine-disulfide exchange (32, 33). Detailed characterization of Rv0081 would therefore lead to the mechanisms by which Mycobacteria sense varying levels of oxygen and switch between dormancy and actively dividing forms.

While the DNA binding property and the repressor activity of Rv0081 are well established, the specific stimulations that regulate binding and dissociation of Rv0081 from DNA, thus influencing its regulatory property, remain unclear. Although ArsR/SmtB family of regulators are known to respond to toxic levels of metals by virtue of the conserved metal binding motif - **ELCV(C/G)D**, termed as the ‘metal binding box’ in α-3 and α-5 helices (22, 34), we and others observed that Rv0081 does not have the conserved metal binding residues either in α-3 or α-5 helices but has DNA binding residues in α-4 region. This suggested that DNA-binding properties of Rv0081 might be metal independent. In Mtb, metal independent DNA binding is regulated by either thiol switch or post transcriptional modifications, such as phosphorylation (35–38), acetylation (39, 40), or methylation (41, 42) and we can postulate one or combination of such mechanisms regulates the Rv0081. The formate hydrogen lyase genes in *E. coli* are regulated by the *fhlA* transcription factor, which appears to sense formate levels in triggering the expression of FHL genes (43, 44). It might be possible that Rv0081 also senses formate levels, instead of oxygen levels, and thereby influences expression of the *rv0081-rv0088* operon. With the emerging significance of Rv0081 in dormancy and our own observation of its indirect dependence on GroEL (45), we have determined the crystal structure of Rv0081and identified the possible stimulations that may regulate binding of Rv0081 to its cognate DNA element. The mechanism of regulation by Rv0081 is proposed, which might be due to post-translational modification of Rv0081 during control of gene expression.

## RESULTS

### Overall structural features of Rv0081

The structure determined by Molecular Replacement showed four chains in asymmetric unit. The best Molecular Replacement solution was obtained only in the space group P4_1_2_1_2. Moreover, this solution did not yield any short contacts between symmetry related molecules thereby suggesting the solution to be correct. The structure was refined to reasonable geometrical parameters (Table 1). One cell length being very long could be explained by the presence of two dimers in the crystal asymmetric unit arranged in a side-to-side manner. Earlier it has been reported that the molecule is likely to be a monomer, however, recently it has been reported that Rv0081 exist in dimeric form (46) and also forms higher oligomers at higher concentration (47). Therefore, we tested the presence of dimeric association through multi-angle light scattering coupled with size exclusion chromatography (SEC-MALS). These results confirmed that *in vitro* purified protein is indeed dimeric in tested buffer conditions (Supplementary Figure S1A-C) and supported earlier finding (46).

**Table 1:**
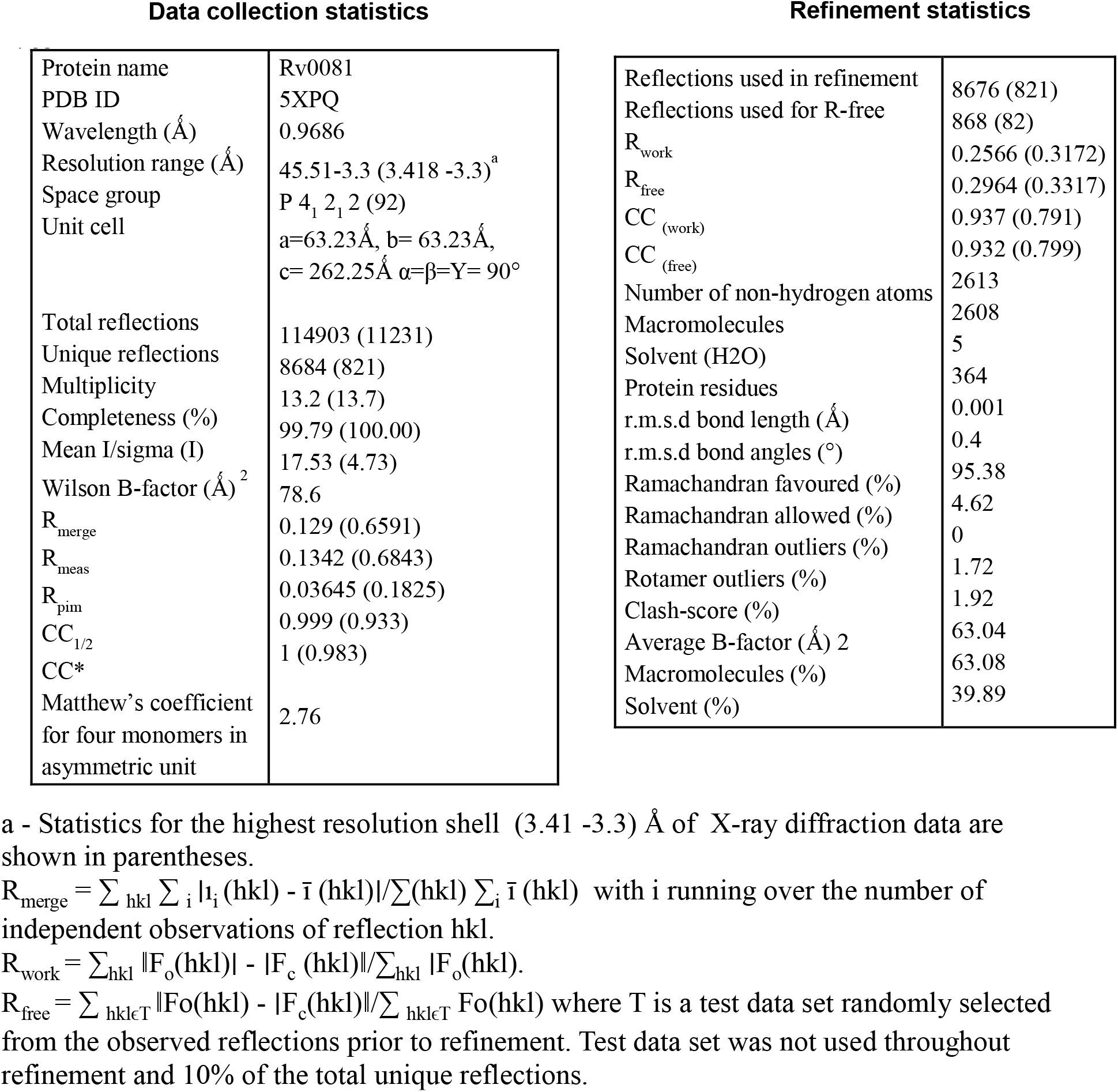
Crystallography data collection and refinement statistics.

Furthermore, to confirm the SEC-MALS results of the biological assembly, we calculated the area buried among the four different Chains. The Chains A and D; and B and C possess interface area of 1499.5, 1465.9 Å^2^ respectively. On the other hand, the interface area between chains AD and chain BC is 609 Å^2^. Chain A and Chain D thereby appear to form one biological dimer whereas Chain B and Chain C form the other biological dimer. The four chains in the asymmetric unit together therefore constitute two biological dimers (Figure 1).

**Figure 1.**
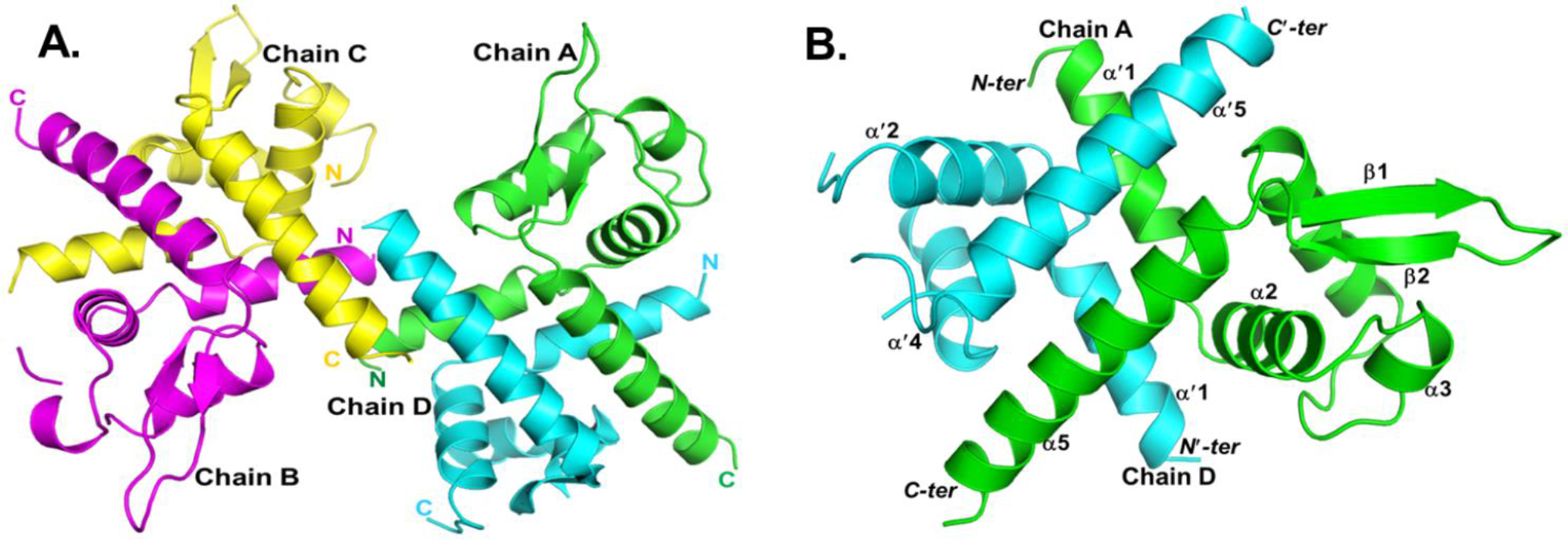
Structure of the Rv0081 dimer. The structure of Rv0081 is typical of the ArsR/SmtB family of metal-dependent transcriptional repressors. (A) Crystal asymmetric unit contains four identical chains as shown in four different colours. The extremities of the polypeptide chains at either of the N- and C- termini make up the interface between two dimers. At this stage it is not clear if the tetrameric assembly is biologically relevant or not. (B) Two monomers possess an extensive monomermonomer interface, clearly suggesting that the biological assembly is that of a dimer. The interface is principally made up of the 1^st^ and the last helices. Only in one chain, i.e., in Chain A, the entire molecule could be fitted in the electron density. In all the other chains, there are disordered regions which could not be fitted in electron density (see text for details).

In each of the chains, a number of residues at the N- and C- termini were not visible in the electron density, and hence were excluded from the final model. Moreover, the region of the structure that is likely to be involved in DNA-binding is highly disordered in all the four chains. The final model therefore constitutes of residues Ser 3 to Ala 102 in Chain A; Glu 4 to Leu 40, Leu 46 to Ser 48, Asn 50 to Ala 102 in Chain B; Glu 4 to Val 44, Glu 47 to Ala 102 in Chain C; and Glu 4 to Val 36, Asn 50 to Ala 65 and Tyr 75 to Val 101 in Chain D. Presumably because of such large segments of structural disorder, the average temperature factor is high, leading to poor diffraction quality of the crystals.

The overall structure is similar to other proteins in the ArsR/SmtB family shown in Supplementary Table S2. Structural superposition with apo-CzrA (PDB: 1R1U) and apo-SmtB (PDB: 1R1T) yielded overall rms deviation of 1.8 Å and 1.9 Å (Supplementary Figure S2A and S2B) respectively. Maximum deviation is seen in the region where DNA-binding site is located. The overall structure is archetypical winged helix-turn-helix motif DNA binding proteins of this family. The disordered DNA binding region of the structure is suggestive of induced fit binding to the DNA.

### Rv0081 lacks metal binding motif but has conserved DNA binding residues

CzrA and SmtB, as indeed all other members of the ArsR/SmtB family transcriptional repressors, are known to bind DNA in a metal-regulated manner. There are three major classes of metal binding regulators in this family, those which bind metal defined by ‘α3 motif (e.g. ArsR), ‘α5 motif (e.g. CzrA, NmtR, and CmtR) and ‘α3 and α5 together (e.g. SmtB, CadC and ZiaR). We compared the structural regions of Rv0081 with the corresponding region of metal bound CzrA-Zn_2_ (PDB: 1R1V) with overall rmsd of 1.5 Å and SmtB-Zn_2_ (PDB: 1R22) with overall rmsd of 1.8Å. Surprisingly, we found that the residues presumably involved in binding metals are mutated in Rv0081 (Figure 2A). For example, in α5 family, in the structure, CzrA-Zn_2_, the residues involved are Asp 84, His 86, His 97 and His 100 whereas in α3-α5 family structure, SmtB-Zn_2_, the residues are Asp 104, His 106, His 117 and Glu 120. The superposed residues in Rv0081 are Ala 79, Asp 81, Val 92 and Arg 95 respectively, with both α5 and α3-α5 family. The superposition of Rv0081 with α3 family proteins was not performed owing to unavailability of the crystal structure. Multiple sequence alignment of Rv0081 with ArsR (α3), CzrA (α5) and SmtB (α3-α5) proteins also showed that Rv0081 did not have metal binding residues (Figure 2B). Further, we examined the Rv0081 structure for potential metal binding sites in the PAR3D program (48) and found that there are no potential metal binding sites in Rv0081. It therefore appears that the activity of Rv0081 binding to DNA might not be regulated by metals.

**Figure 2.**
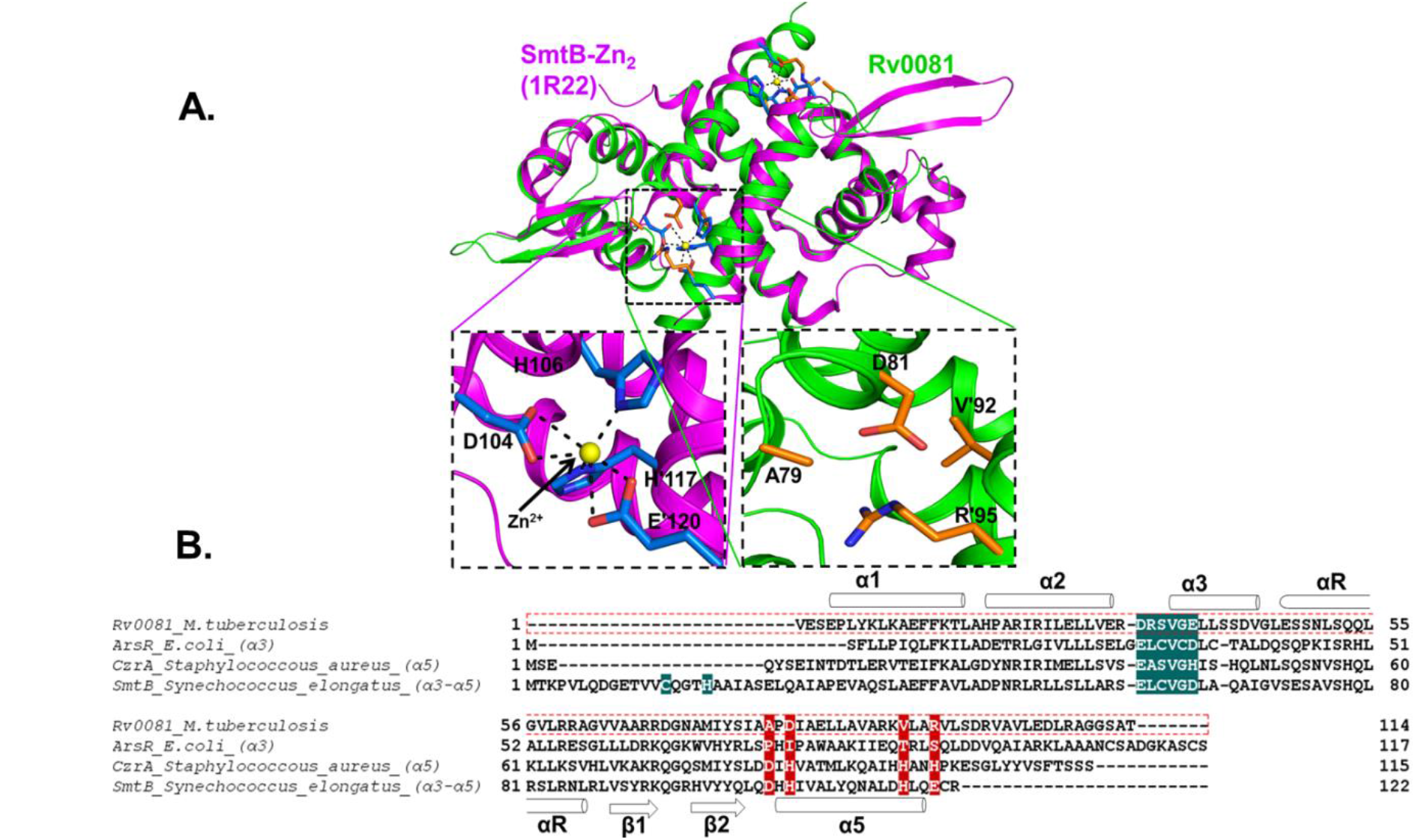
Superposition of Rv0081 structure with the SmtB-Zn2 structure bound to metals (PDB: 1R22) and multiple sequence alignment (MSA) of representative of α3, α5 and α3-α5 regions of ArsR/SmtB family with Rv0081. (A) In the α3-α5 family, SmtB-Zn_2_ complex (purple ribbon structure), Zn^2+^ ion only seen at α5 region, not at α3 region. Residues of SmtB, which are coordinated with Zn^2+^ are Asp 104, His 106, His’ 117 and Glu’ 120 at α5 region, where the latter two residues are contributed by another monomer in the dimer. Structural superposition shows that these residues in the corresponding region of the Rv0081 structure (green ribbon structure) are Ala 79, Asp 81, Val’ 92 and Arg’ 95. Therefore, the metal coordinating ability of Rv0081 appears to have been lost during evolution. Similar comparisons with the regions of α3 and α5 also reinforce the same conclusions. (B) Multiple sequence alignments (MSA) of ArsR (α3), CzrA (α5) and SmtB (α3-α5) protein sequence with Rv0081sequence, the region highlighted with green and red represents α3 and α5 regions respectively. Superpositions and MSA of α3, α5 and α3-α5 regions of ArsR/SmtB family with Rv0081 showed that Rv0081 might have lost its ability to bind metals.

By comparing the Rv0081 structure with other ArsR/SmtB transcription factors, we could delineate the regions likely to play important roles in DNA recognition (Figure 3A). From the structural superposition of NolR-DNA complex of *Sinorhizobium fredii* (PDB: 4OMY) there are two kinds of residues that are seen to interact with the DNA- those which are involved in contacting the bases of the DNA, and those which interact with the phosphate backbone. In the NolR-DNA complex structure the base recognizing residues are Gln 56, Ser 57, Ser 60 and Gln 61 (49). Superposition of the two structures suggests that these residues in Rv0081 would be Ser 48, Ser 49, Ser 52 and Gln 53. Similarly, the phosphate binding residues in Rv0081 would be Lys 15, His 19, Arg 22, Gln 54, Asn 71 and Tyr 75. In the surface diagram representation of dimer of Rv0081 in Figure 3B, base recognizing residues denoted as yellow lie at the extremity of the dimer and phosphate binding residues in red color span the connecting the entire surface. Thus, residues involved in DNA recognition could be predicted from comparisons with other structures that are available in complex with cognate DNA.

**Figure 3.**
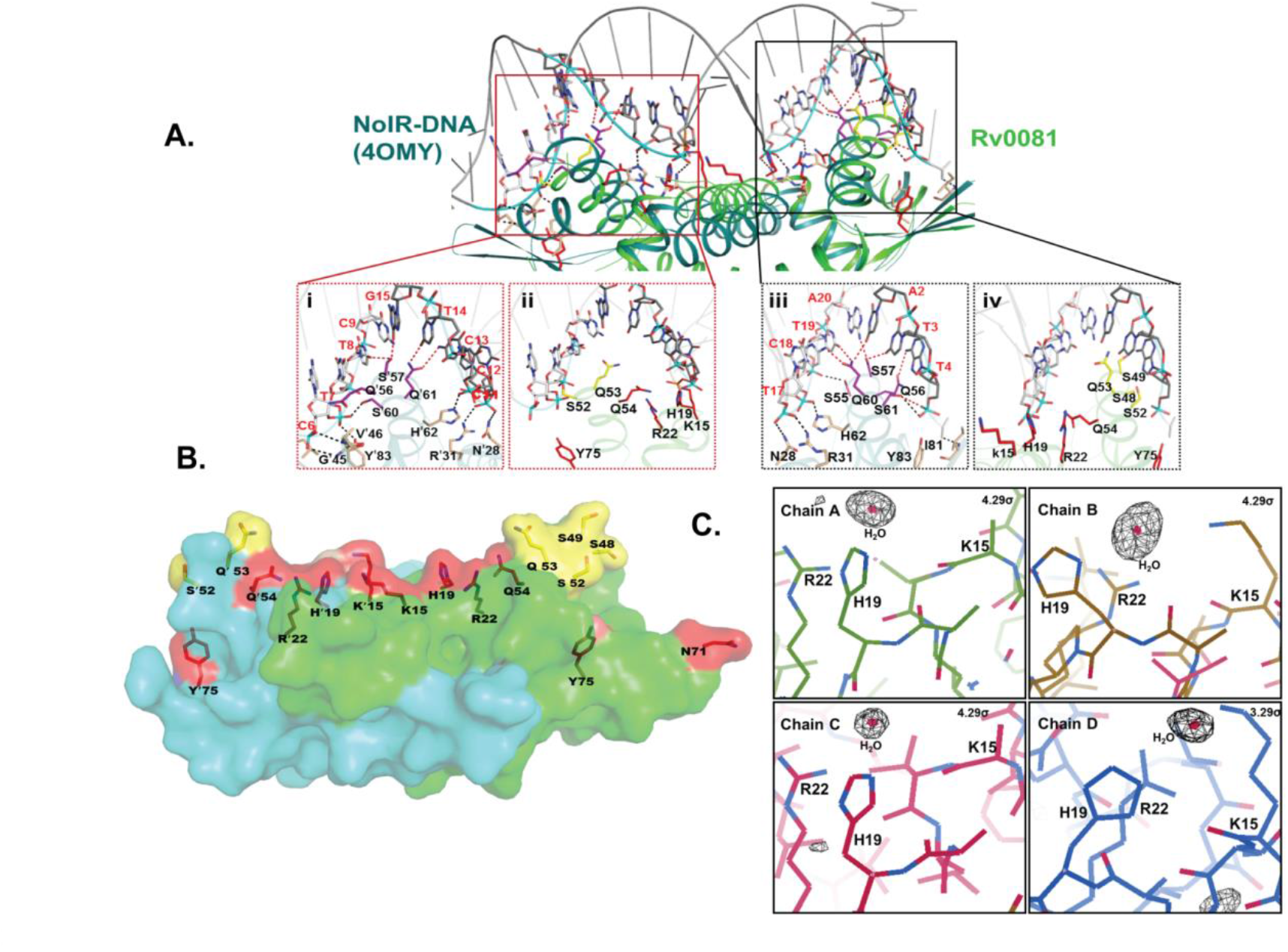
Superposition of Rv0081 structure with the NolR-DNA structure bound to DNA. (A) The superposition of Rv0081 with NolR (PDB: 4OMY) clearly indicates two different kinds of binding modes – (1) residues which are involved in base-recognition (red dotted lines), and (2) those involved in contacting the phosphate backbone (black dotted lines). DNA binding domains of NolR are highlighted in box. NolR residues interacting with bases are highlighted in purple and those with phosphate in light brown color. Bases of DNA are highlighted in white and gray and phosphate backbone in cyan color (Box i and iii). The superposed residues of Rv0081 that might potentially interact with bases are highlighted in yellow and those with phosphate in red color (Box ii and iv). Thus, the superposition suggests the sequence of the DNA involved in DNA-Rv0081 interactions. (B) Surface diagram of dimer of (chain A, green and chain D, cyan) Rv0081 Structure shows that the base-contacting residues (yellow) lie at the extremities of the structure, whereas the phosphate contacting residues (red) fill in the intervening surface of the structure. (C) Clustering of basic residues in Rv0081, i.e., Lys 15, His 19 and Arg 22, is suggestive of anion binding site. Fo-Fc map contoured at 5 level shows a strong density in this region. However, due to limited resolution of the structure we have avoided ascribing this density to any anion. See text for details.

In the Rv0081 structure, Lys 15, His 19 and Arg 22 form a cluster of basic residues, suggesting their possibility of anion binding. Indeed, Fo-Fc maps show a strong peak at 5 above mean electron density level in three of the four chains (Figure 3C). We have currently modeled a water molecule as the low resolution structure does not permit us to correctly identity of ligand in this strong density. However, such a clustering of basic residues is reminiscent of anion binding sites such as sulfates (50, 51) and phosphates (52, 53), or acetate or formate (54, 55). A higher resolution of Rv0081 structure will likely reveal the correct identity of anion binding at these sites.

### DNA-binding by Rv0081

#### a. Interaction studies on Rv0081 protein with auto-regulatory elements

DNA-binding of Rv0081 was confirmed for a 33 bp oligonucleotide with its sequence derived from upstream sequence of the Rv0081- Rv0088 operon. This region encompasses the R1 and R2 regions, where the protein is believed to bind first to the R2 region, and then cooperatively to the R1 region (23). From the structure we identified five bases which are likely to be contacted by the protein. These analyses suggested that Rv0081 might bind to the sequence <u>**TATCT**</u> in the R1 region and <u>**TCTTC**</u> in the R2 region (Figure 4A). We mutated these five bases in one or both of the R1 and R2 regions into <u>**GGGGG**</u> and tested binding by the protein (Figure 4A). Electrophoretic mobility shift assay (EMSA) analyses with the wild-type or mutated oligo’s are shown in Figure 4B. As expected, the protein binds to the native sequence (Figure 4B). This binding is sequence specific since bound biotinylated Wt-oligos competed with non-biotinylated Wt-oligos but did not compete with scrambled-oligos RS1, RS2 and RS3 (Figure S3A). EMSA results clearly show that Rv0081 binding affinity diminishes in the R1 mutated oligo (M2), whereas it is almost completely abolished in the R2 mutated (M1) region (Figure 4B). Corroboratively, surface plasmon resonance (SPR) with purified Rv0081 and immobilized oligos [Wt, M1 (R2 mutant) and M2 (R1 mutant)] shows reduced binding for the R1 mutated region (M2) and almost completely abolished binding for R2 mutated (M1) region (Figure 4C). Thus, these results indicate that the Rv0081 protein binds to the R2 region more efficiently than the R1 region, confirming the earlier findings (23).

**Figure 4.**
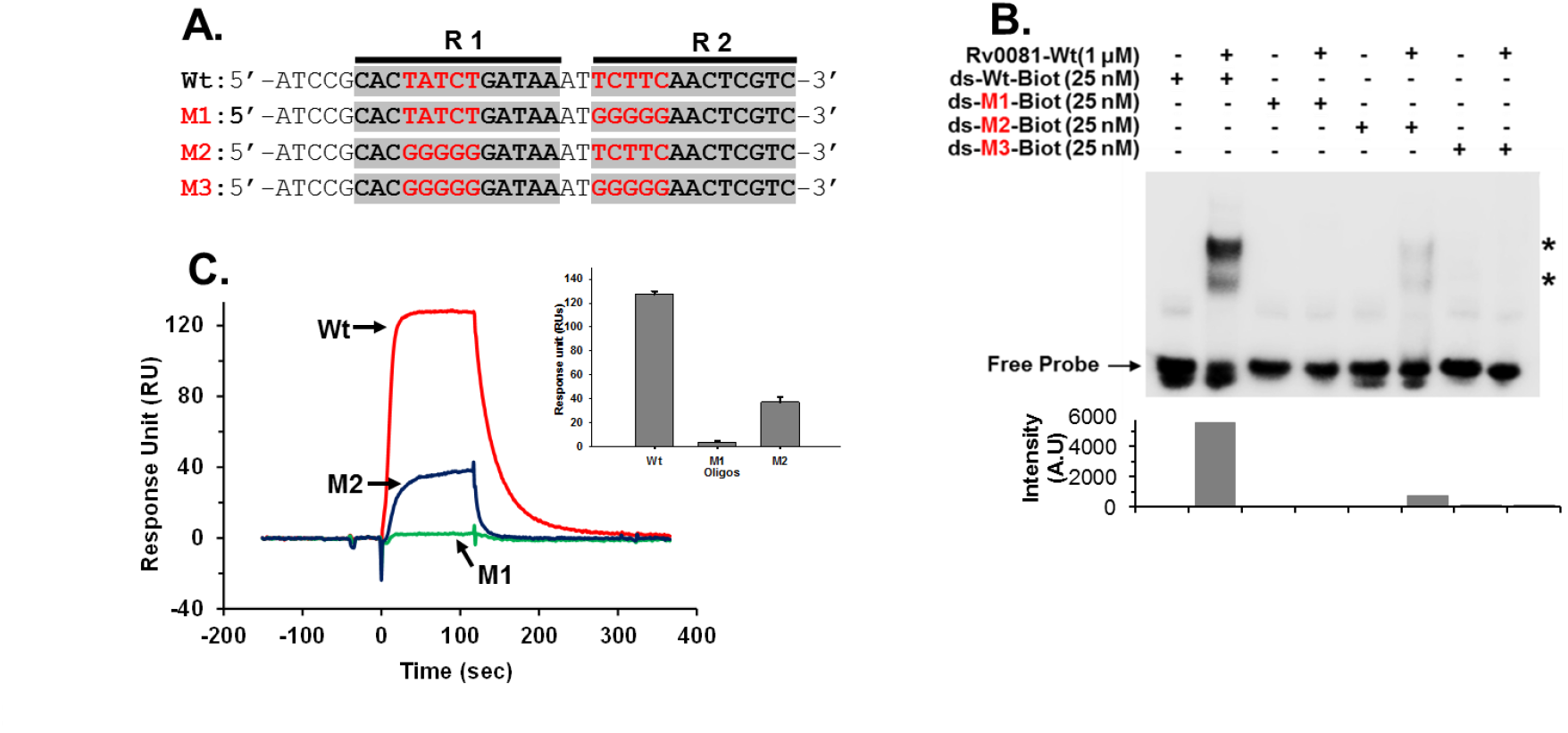
Electrophoretic mobility shift assays (EMSA) and surface plasmon resonance (SPR) of the wild-type Rv0081 with different oligo sequences. EMSA and SPR were carried out as described in the text with a 33 base pair native region (Wt) and mutated oligos M1, M2 and M3. (A) Two DNA motifs interacting with Rv0081 are labelled as R1 and R2. Five potential bases both in R1 (TATCT) and R2 (TCTTC), interacting with Rv0081 in (Wt) oligo were mutated to GGGGG in either R1 (M2) or R2 (M1) as well as both (M3). (B) EMSA was performed with the Rv0081 protein (1μM) and biotinylated DNA probes (25 nM) Wt, M1, M2 and M3. Shifted bands are indicated by ‘*’ and the bar plot below shows the intensity of shifted bands. (C) SPR analysis of the Rv0081 protein (100 nM) with immobilized DNA probes of either Wt (red), M1 (green) or M2 (blue) mutants performed using Biacore 2000. Average response units for the Rv0081 protein with ds-Wt as well as M1 and M2 oligos are shown using a bar graph inset (Data are mean + SD of three? experiments). Weak binding to M2 (R1 mutant) and no binding to M1 (R2 mutant) oligo sequences suggest that the Rv0081 protein might initially bind the R2 region, and then bind cooperatively to the R1 region.

#### b. Mutational analysis of Rv0081 suggest modulation of DNA binding

Based on the superimposition of Rv0081 structure with NolR-DNA complex, we identified four key residues of Rv0081 for interaction with DNA bases. These residues S48, S49, S52 and Q53 were selected for the mutational analysis. We generated two mutants of Rv0081 proteins– one where all these four residues were mutated to Ala (Rv0081-S/Q-A), and the other where Ser were mutated to Asp (Rv0081-S-D; Figure 5A) – and EMSA as well as SPR experiments were performed to compare the interaction of Rv0081 and its mutants (Rv0081-S/Q-A, Rv0081-S-D). Binding of Rv0081 and Rv0081-S/Q-A mutant were in concentration dependent manner with ds-Wt DNA (EMSA). The binding was reduced in Rv0081-S/Q-A mutant whereas it was almost completely abolished in Rv0081-S-D mutant (Figure 5B). Similar pattern of interactions was observed with both the mutant proteins using the SPR. Rv0081-S/Q-A mutant showed moderate binding response, whereas Rv0081-SD displayed substantially reduced binding response. It is important to point out here that Rv0081-S/Q-A failed to show equivalent binding response to that of Rv0081 even at a concentration that is 32-times (3200 nM) than that of Rv0081, whereas Rv0081-SD showed barely detectable binding response 6400 nM concentration (Figure 5C, Figure S4A).

**Figure 5.**
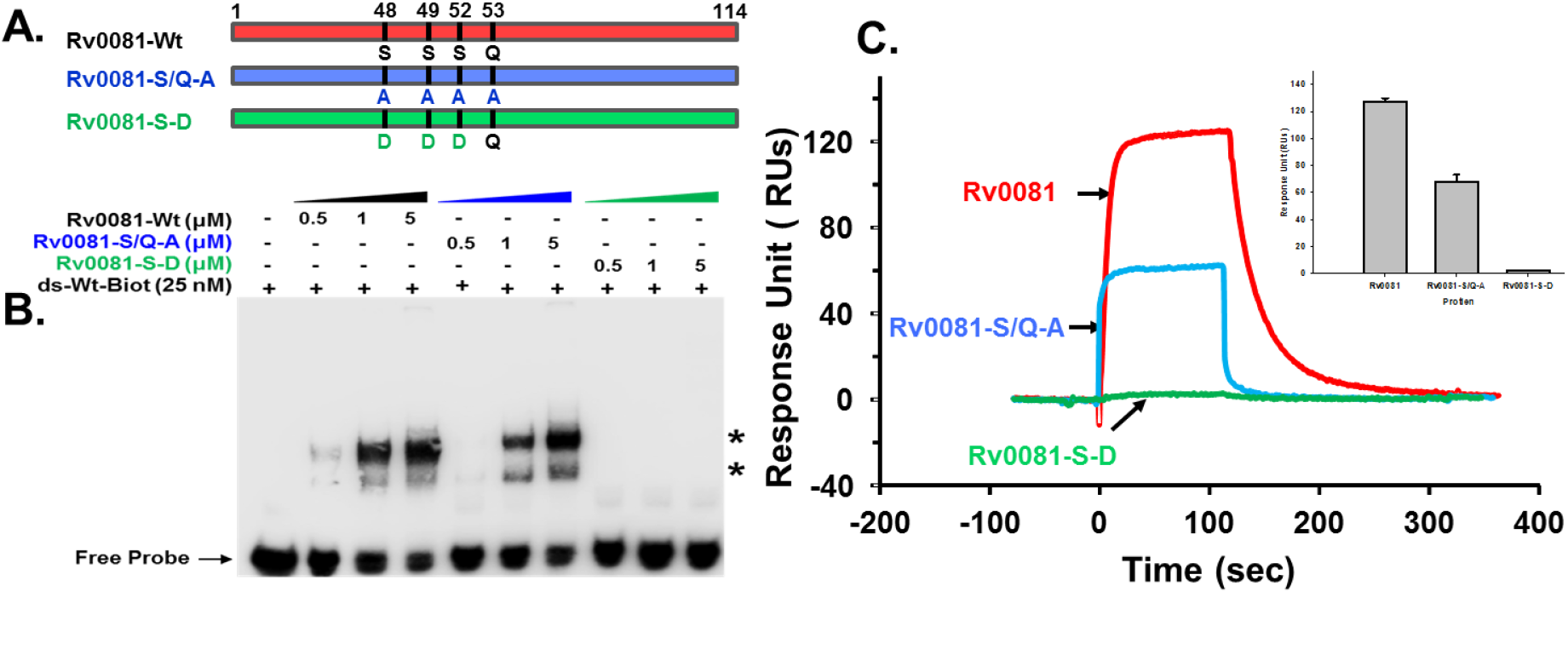
Electrophoretic mobility shift assays (EMSA) and surface plasmon resonance (SPR) of the wild-type and mutants (S/Q-A, S-D) of Rv0081 with wild-type oligonucleotide sequences. (A) Schematic representation of the amino acid residues of wild-type Rv0081 protein (Rv0081-Wt) and its mutants Rv0081 (S/Q-A) and Rv0081 S-D t. (B) EMSA was performed with increasing concentration (0.5, 1and 5 μM) of either Rv0081-Wt, Rv0081-S/Q-A or Rv0081-S-D proteins with wild-type biotinylated DNA probe (Wt) and shifted bands are indicated by ‘*’. (C) Binding interaction of the Rv0081 or its mutant proteins with the wild-type oligonucleotide sequence. The biotin-labelled wild-typeoligo (test flow cell) or the scrambled oligo (control flow cell) was immobilized on a streptavidin chip. The specific binding response (shown as sensogram overlay plots) of Rv0081 (100nM), Rv0081-S/Q-A mutant (3200nM) and Rv0081 S-D mutant (6400nM) to the wild-type oligonucleotide is shown in red, blue and green, respectively. The inset bar graph shows the average binding of all the three proteins to ds-Wt oligos (Data are mean + SD of three? experiments).

#### c. Kinetics of Rv0081-DNA binding

To calculate the affinities of the Rv0081 and its mutants to the wild-type or mutant oligo’s, different concentration of proteins were flown over the oligo-immobilized chip. The SPR results clearly showed that the native oligonucleotide sequence (ds-Wt DNA) binds with greater affinity to the native Rv0081 protein. In order obtain the affinity for this interaction, the data was fitted with a Bivalent model (BIAevaluation) (χ^2^=2.74). The analyses indicated *K*_D1_ = 1.04 x 10^-7^ M and *K*_D2_ = 5.2×10^-2^ M (Figure 6A, Table 2), suggesting that one of the binding sites has higher affinity. This is consistent with our EMSA and above SPR analysis data (Figure 4B-C), confirming that R2 region has the higher affinity for Rv0081. The R2 mutated oligonucleotide (M1 oligo) failed almost completely to bind (Figure 4C), on the other hand, the R1 mutated oligonucleotide (M2 oligo) bound to Rv0081 with 100-fold lower affinity (*K*_D_ 1.56×10^-5^ M; Figure 6B, Table 2.). Effect of mutation on low affinity R1 region also suggests the possibility of cooperative binding of both R1 and R2 regions with Rv0081.

**Figure 6.**
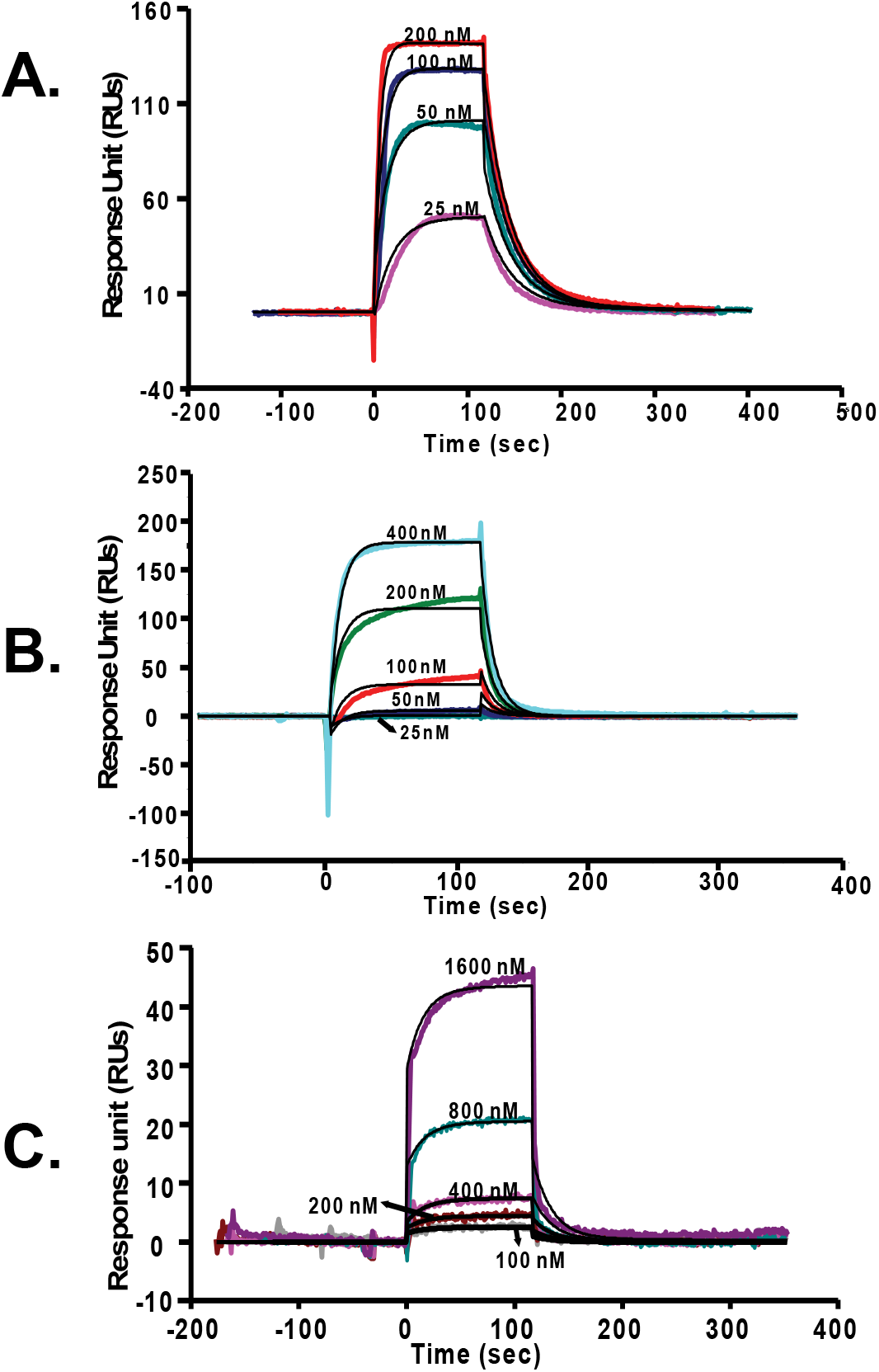
Binding analysis of Rv0081 (Wt) or the mutant protein (Rv0081-S/Q-A) to the immobilized oligos (Wt/M2 oligos). The sensogram overlay plots of Rv0081 (Wt protein) flowed at different concentrations over the wild-type (A) or the mutant oligo M2 (B) is shown. Similarly the sensogram overlay plots of the mutant protein (Rv0081-S/Q-A) is shown in (C). The solid lines represent the fitting of the data either by bivalent model (A, C) or with the 1:1 Langmuir binding model. The details of kinetics are described in Table 2.

**Table 2:** Kinetic and affinity data for interactions of Rv0081 and its Rv0081-S/Q-A mutants with its auto-regulatingelements (wt oligo) and its R1 mutant (M2 oligo)

| Oligo | Protein | $k_{d1}$ (1/s) / $k_{a1}$ (1/Ms) | $K_{D1}$ (M) | $k_{d2}$ (1/s) / $k_{a2}$ (1/Ms) | $K_{D2}$ (M) | Rmax | Chi<sup>2</sup> | Fitting model |
| --- | --- | --- | --- | --- | --- | --- | --- | --- |
| Wt | Rv0081 | 3.41x10<sup>-2</sup>/3.28x10<sup>5</sup> | 1.04x10<sup>-7</sup> | 26.101/4.97 | 5.25x10<sup>-2</sup> | 161 | 2.74 | Bivalent |
| Wt | Rv0081-S/Q-A | 5.54x10<sup>-2</sup>/2.82x10<sup>3</sup> | 1.96x10<sup>-5</sup> | 0.261/4.38 | 2.17x10<sup>-4</sup> | 101 | 0.549 | Bivalent |
| | | $k_d$ (1/s) / $k_a$ (1/Ms) | $K_D$ (M) | | | | | |
| M2 | Rv0081 | 9.92x10<sup>-2</sup>/6.34x10<sup>4</sup> | 1.56x10<sup>-6</sup> |  | - | 784 | 13.5 | Langmuir (1:1) |

$k_a$ , association rate constant; $k_d$ , dissociation rate constant; $K_D$ , equilibrium dissociation constant. Data shown here are representative of three independent experiments.

**Table 3:**
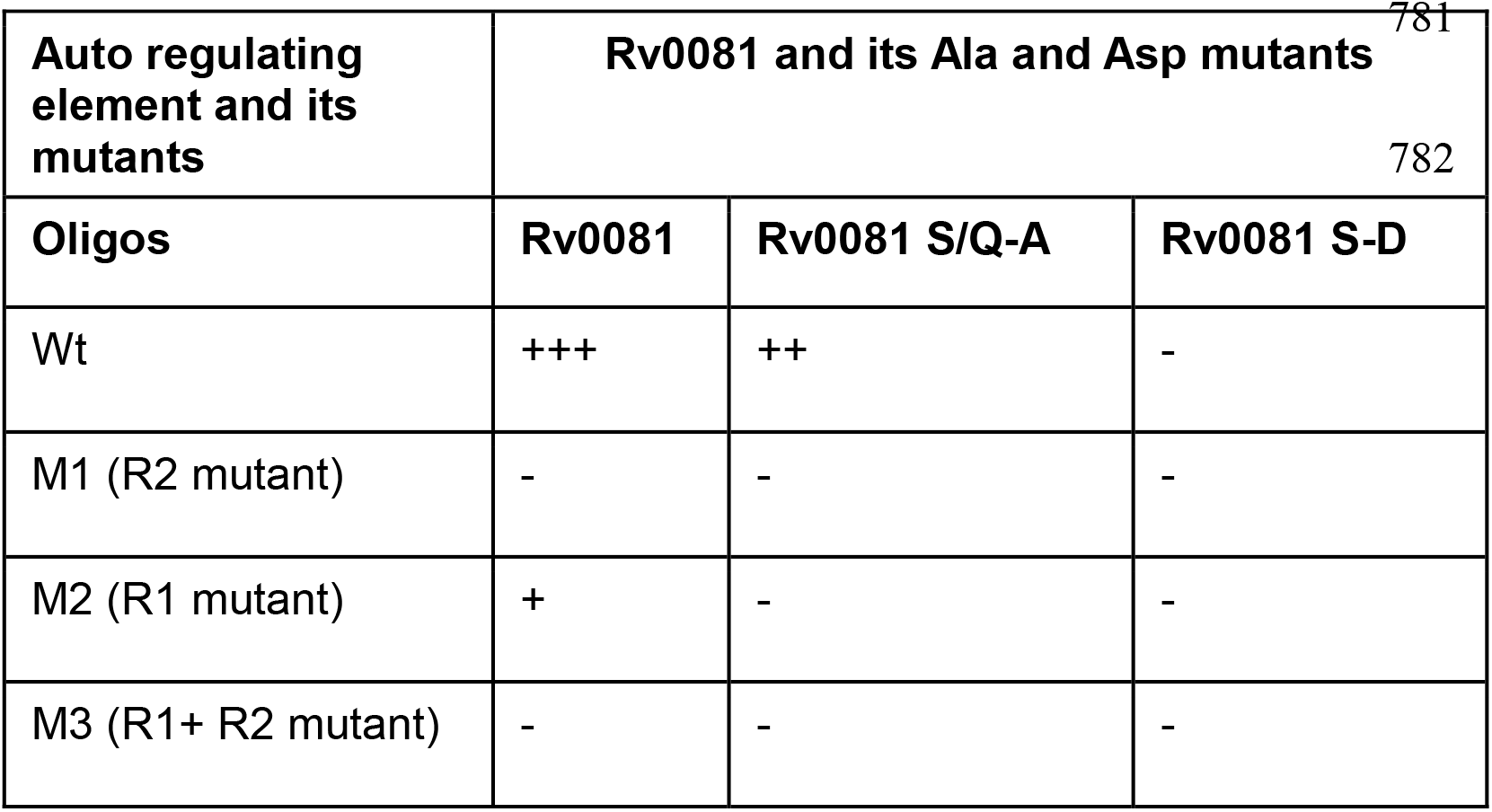
Summary of interactions of Rv0081, its Ala and Asp mutants with Wt-type oligo R1, R 2 and both R1 and R2 mutants.

### Rv0081 is phosphorylated in hypoxia

The presence of multiple Serine residues in the DNA binding region of Rv0081 protein is suggestive of post-translational modification of these for regulating DNA binding. We therefore hypothesized that in absence of metal-binding, the DNA binding activity of Rv0081 might be regulated by post-translational modifications – i.e., phosphorylation of Serines, rather than by metal ions. In any case, the structure does not show presence of putative metal binding site (48). We created mutants of Rv0081, where the Ser-residues are mutated to Asp. The Ser to Asp mutants were created to mimic phosphorylation of Rv0081, where we could test the hypothesis that regulation of DNA binding activity was mediated by post-translational modification. As described above, the Ser to Ala mutants bind DNA with lower affinity (*K*_D1_= 1.96 x 10^-5^ M, *K*_D2_= 2.17 x 10^-4^ M, Figure 6C, Table 2), whereas we expected that the Ser to Asp mutants might abolish binding completely. Consistent with this hypothesis, the results on these EMSA and SPR experiments (Figure 5) show clearly that the Ser to Asp mutants fail to bind DNA.

Finally, to test if the protein was really phosphorylated under normoxic and hypoxic conditions, we probed the protein from crude cell lysates with anti-phospho-Ser antibodies. Cells grown under these two conditions showed that the expression of Rv0081 did not change much under the two conditions. However, under the hypoxic condition, there was a marked enhancement in the phosphorylation of the protein (Figure 7, Figure S5). This experiment thus, confirmed that Rv0081 might be differentially phosphorylated under these two environmental conditions, and that its DNA-binding activity might be regulated by such a post-translational modification.

**Figure 7.**
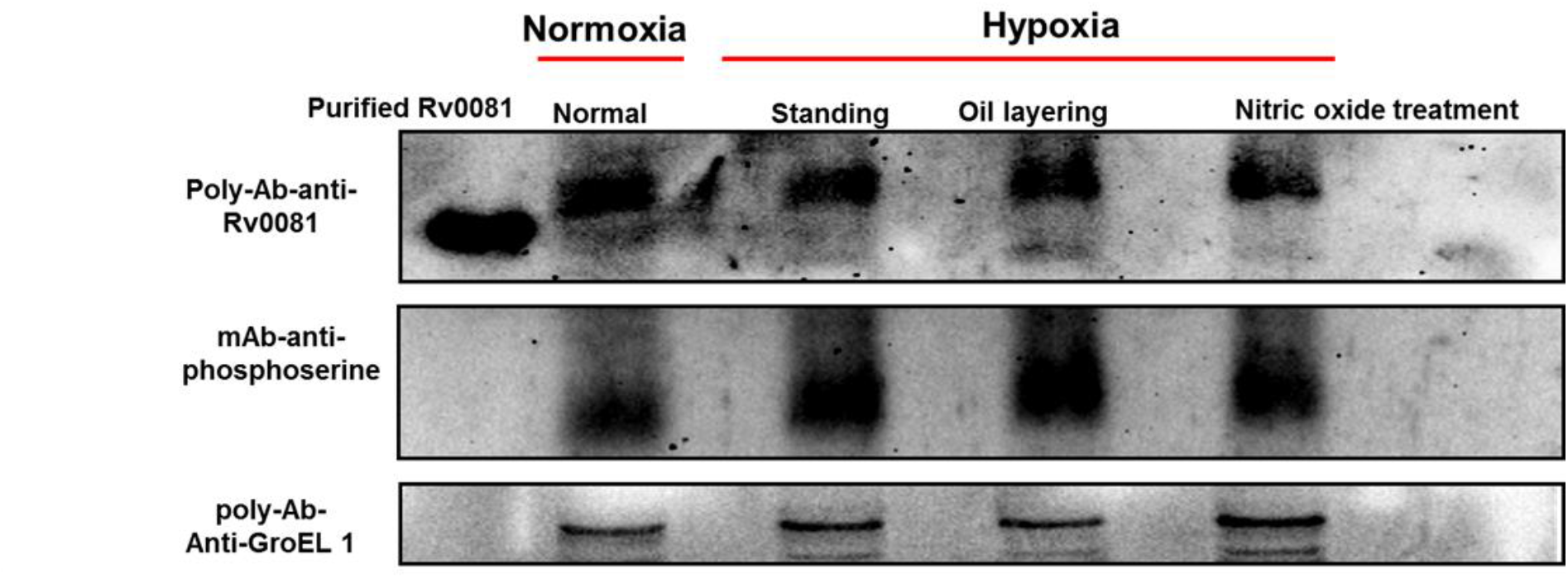
Phosphorylation of Rv0081 under Hypoxia. Levels of phosphorylation of Rv0081 under normal and hypoxia induced cultures of *M. tuberculosis* were measured using Co-IP method. *M. tuberculosis* cultures were induced with hypoxia by either keeping the culture with low aeration (LA described in details in material methods) low aeration with overlaid mineral oil (MO) or low aeration with 250 μM Nitric oxide (NO) for 1 day Cell lysates were prepared in lysis buffer and resolved on denaturing SDS-PAGE. Proteins were blotted on PVDF membrane and the mAb-anti-phosphoserine and then after with polyAb-anti-Rv0081. Purified Rv0081 (expressed in *E. coli*) was used as a control. Result suggested a marked enhancement in the phosphorylation of the protein under hypoxia condition as compared to normal, although expression of Rv0081 did not change.

## DISCUSSION

Although the Rv0081 crystals diffracted poorly, we could successfully determine its structure to 3.3 Å resolution. The dimeric assembly of Rv0081 is canonical of transcriptional repressors of ArsR/SmtB family. In the crystal structure, the DNA binding regions are ordered only in one of the four chains. However, they are juxtaposed symmetrically in the biological dimer suggesting that the mode of DNA binding is similar to that of the most bacterial transcriptional repressors of this family (22). The DNA binding by Rv0081 can be imagined as being spread through one surface of the molecule (Figure 3B), with the base recognizing residues lie at the extremities of the dimer, whereas the entire length of the intervening surface being dotted with basic amino acids which are likely to interact with the phosphate backbone of the DNA. The phosphate-recognition residues from the two monomers therefore cover the entire surface of the dimer between the two base-recognition sites.

Whereas the DNA binding region was easy to identify, the metal binding region of the protein was not easily discernible. He *et al*. have reported that Rv0081 fails to bind metals (23) In the other structures of the ArsR/SmtB family of proteins, one or two metal binding sites are known, which are designated as either α3, α5 or both α3-α5 (22, 34). Superposition of the Rv0081 structure with other family members of the ArsR/SmtB revealed that in both these putative metal binding regions the residues in Rv0081 are either non-polar, or mutated as to be devoid of metal-binding. Further, multiple sequence alignments and phylogenic analysis of different classes (α3, α5 and α3-α5) of ArsR/SmtB family with Rv0081 amino acid sequence also supported our observation (Supplementary Figure S6, S7 S8, S9). We therefore agree with the reported study that Rv0081 is unlikely to bind cations (23). As other members of this family sense metal dependent redox changes, the mechanism of Rv0081 to sense the hypoxic conditions clearly appears to be distinctly different and metal-independent. In absence of cation binding, the mode of transcriptional regulation in response to hypoxic conditions therefore remains enigmatic. Moreover, in absence of metal-binding, the regulation of DNA binding and the accompanying allostery of Rv0081 are not clearly understood.

We hypothesized that under hypoxic conditions, the protein might get post-translationally modified, e.g., phosphorylated, and thereby alter its DNA-binding ability. The three Serines which are postulated to be involved in base-recognition become excellent candidates to test this hypothesis (Figure 5). We mutated the four residues Ser 48, Ser 49, Ser 52 and Gln 53 to Ala, and the three residues Ser 48, Ser 49 and Ser 52 to Asp. The two mutants of protein show significant differences in DNA binding ability. Whereas the former binds DNA with lower affinity, the latter almost failed to bind to DNA (Figure 5C). The Serine to Aspartate change being a mimic of a phospho-Serine, this result is suggestive of Ser-phosphorylation in Mtb Rv0081. Indeed, Mtb cultures grown under normoxic and hypoxic conditions clearly exhibit differential phosphorylation of the Rv0081 protein, with elevated levels of phosphorylation under the hypoxic conditions. Thus, in absence of metal-binding, this member of the ArsR/SmtB family transcriptional repressor appears to have evolved a novel mechanism for sensing hypoxic conditions.

The oxygen-sensing mechanisms in most transcriptional repressors, apart from reversible metal-binding, are mediated by thiol redox switch. Crystal structures of many members of this family have revealed these mechanisms in details (48–51). Yet, Rv0081 does not possess any such canonical site for sensing oxygen levels. It could be hypothesized that, Rv0081 is being the first gene on the formate hydrogenlyase operon, and acting as a repressor of this operon, might therefore also be activated by formate ions. This hypothesis will need to be explored further.

Finally, the structure of Rv0081 and accompanying solutions studies reveal that different levels of DNA-binding regulations by Rv0081 might exist in *M. tuberculosis*. On one hand, post-translational modification diminishes DNA-binding; on the other hand there is an untested hypothesis that it might be regulated by formate ions. Galaghan *et al*. have reported Rv0081 to be a regulatory hub of many genes in Mtb (20), and it is likely that such multiple mechanisms of DNA-binding offer redundancy to switch-off and -on different genes via different mechanisms. The crystal structure of Rv0081 therefore reveals mechanisms of DNA-binding which are distinct from other members of the ArsR/SmtB family members.

## MATERIALS AND METHODS

### Purification, Crystallization and Structure Determination of Rv0081

The (His)_6_-tagged Rv0081 protein was purified to homogeneity (Supplementary Methods) and used for crystallizations. Prior to setting up crystallizations the protein was centrifuged at 13800 x g for 20 min at 4 °C and concentrated to 1.5-2.0 mg/ml. Diffractable crystals were obtained in a condition with 0.1M CH_3_COONa, 0.2-0.4 M (NH_4_)_2_SO_4_ and 25% w/v PEG 4000 in 2 µL drops set at room temperature. The crystals were stabilized in crystallization condition with 20% glycerol before flash-freezing in liquid-nitrogen for data collection at 100K.

To determine the structure of Rv0081 using molecular replacement six different models from Protein Data Bank with PDB-IDs 4GGG, 2KJB, 3PQK, 1R1U, 2KKO and 3GW2 were used. Initial phasing was done using dimer of these models using Phaser (62). Initial refinement was done in CCP4i in REFMAC5 (63) module and visualized in COOT (64). Further final manual model building and refinement was done in COOT and PHENIX (65, 66) refinement modules respectively.

### Site directed mutagenesis (SDM)

DNA-binding residues of the Rv0081, namely S48, S49, S52 and Q53, identified by the superimposition of Rv0081 structure with NolR-DNA complex, were selected for mutagenesis. Two mutants were generated where these residues were either changed to alanine or aspartic acid. To generate a mutant of Rv0081 with S48A, S49A S52A, and Q53A (named as Rv0081 S/Q-A henceforth) Forward primer: 5’-TCCTCGGACGTCGGCCTGGAG<u>**G**CG**G**CC</u>AACCTG<u>**G**CC**GC**G</u>CAGCTGGGTGT GCTACGCCGG-3’ and Reverse primer: 5’-CCGGCGTAGCACACCCAGCTG<u>C**GC**GG**C**</u>CAGGTT<u>GG**C**CG**C**</u>CTCCAGGCCGA CGTCCGAGGA-3’ were used. For a mutant of Rv0081 with S48D, S49D and S52D (named as Rv0081 S-D henceforth) Forward primer: 5’- CCTCGGACGTCGGCCTGGAG<u>**GATGAC**</u>AACCTG<u>**GA**C</u>CAGCAGCTGGGTGTG CTACG-3’ and Reverse primer: 5’-CGTAGCACACCCAGCTGCTG<u>G**TC**</u>CAGGTT<u>**GTCATC**</u>CTCCAGGCCGACGTC CGAGG-3’ were used. A PCR reaction containing 0.5 pmol of forward and reverse primers and approximately, 40-50 ng of plasmid (pTZ1070) encoding *M. tuberculosis* Rv0081 gene as the template was carried out using Phusion polymerase (Thermo). Amplified products were digested with approximately 10-15 units of *Dpn*I (NEB) overnight at 37 °C. Digested products were transformed to *E. coli* DH5α strain. Plasmids were isolated from positive colonies (Qiagen) and confirmed by sequencing for the intended mutations.

### Expressions and Purifications of Ala and Asp mutants of Rv0081

The Rv0081 S/Q-A and Rv0081 S-D mutants were expressed and purified in the same way as that of wild-type Rv0081 described above. Purity of mutant proteins was analyzed on 15% SDS PAGE. Proteins were concentrated and snap frozen in liquid nitrogen and stored at −80 °C until further use.

### Electrophoretic mobility shift assay (EMSA)

***DNA probes***: Biotin-labelled and unlabelled oligonucleotides used for performing EMSA (forward and complementary sequence table S2) were synthesized and desalted or PAGE purified (IDT, USA). For double stranded probes, equimolar mixtures of both forward and complementary primers were heated at 95 °C and cooled slowly. Annealed probes were aliquoted and stored at −20 °C until further use. To obtain a mutant DNA probe, Rv0081 crystal structure was superimposed with the NolR from *Sinorhizobium fredii* complexed with dsDNA (PDB: 4OMY) (49). The base pairs to be mutated in mutant oligonucleotides were decided based on the amino acids in Rv0081 which would appear to interact with the bases of DNA, i.e. ATCCGCAC<u>**TATCT**</u>GATAAAT<u>**TCTTC**</u>AACTCGTC, where the underlined bases were mutated into ‘G’.

EMSA were performed using LightShift™ Chemiluminescent EMSA Kit (Thermo) as per the manufacturer’s recommendations. In brief, to test binding of Rv0081 protein (Rv0081-Wt) to 5’-biotinylated ds DNA probe (ds-Wt-Biot), 25 nM of oligoes were mixed with varying concentration of purified Rv0081-Wt protein (0, 0.5, 1.0, 3.0 and 5.0 µM) in reaction buffer [binding buffer (10 mM Tris, 62 mM KCl, 1 mM DTT; pH 7.5), poly-dIdC (50 ng/µl), 2.5% glycerol and 6 mM MgCl_2_] and incubated at RT for 20 minutes. The DNA-protein complexes were resolved on 6.5% non-denaturing-PAGE and transferred to Nylon-66 membrane (GE healthcare). Membranes were crosslinked using UV-trans-illuminator operated at 254 nm for 10 min and 356 nm for 5 min. Signals were detected with streptavidin-HRP-conjugate and Luminol-Enhancer mix as per manufacturer’s instructions. Images were recorded using Imager 600 (GE-healthcare). To test the effect of specific mutations of Rv0081 on DNA binding, purified Rv0081-S/Q-A and Rv0081-S-D mutant proteins were mixed with varying concentrations (0, 0.5, 1.0 and 5.0 µM) in same binding conditions as described before. After 20 min. incubation at RT the complexes were resolved and detected. To test the specificity of DNA on Rv0081 binding, 25 nM of one of the mutant oligo’s (ds-M1-Biot, ds-M2-Biot, ds-M3-Biot) were mixed with 1 µM of either Rv0081-wt or Rv0081-S/Q-A or Rv0081-S-D in reaction buffer and incubated for RT for 20 mins. Complexes were resolved and detected as described. Further, to perform competition assay, 50x molar excess (1.25 µM) of unlabelled oligos were mixed with Rv0081-wt (1 µM) or Rv0081-S/Q-A (1 µM) and mixtures were incubated for 20 min at 4°C. Later, 25 nM of labelled probe (ds-Wt-Biot) was added to the reaction buffer. Complexes were incubated for 10 min at RT and detected as described above. All above experiments were performed in duplicates.

### Surface-Plasmon resonance (SPR)

#### DNA and Protein and binding analysis

To validate the interaction of Rv0081 proteins (Rv0081-wt or Rv0081-S/Q-A or Rv0081-S-D) with ds DNA oligos (biotin-labelled Wt, M1 and M2), surface-plasmon resonance analyses were carried out using the Biacore 2000 system (Biacore AB, Uppsala, Sweden). A sensor chip (GE Healthcare) containing a streptavidin coated surface was activated by three consecutive injections of 1M NaCl and 50mM NaOH. Firstly, to compare the interaction of proteins (Rv0081-wt, Rv0081-S/Q-A or Rv0081-S-D) with ds DNA (biotin-labelled Wt, M1 and M2 oligos), approximately 10 μg/ml of biotin-labelled scrambled dsDNA (RS1), Wt, M1 (R2 mutant) and M2 (R1 mutant) oligos were flown over flow cell-1 (FC-1), FC-2, FC-3 and FC-4, respectively. The immobilization levels on these flow cells were 121 RUs, 120 RUs, 118 RUs and 132 RUs, respectively. Flow cell 1 immobilized with scrambled oligo-RS1 (sequence given in Table S1) was considered as a control. On second sensor chip RS1, Wt, M1 and M2 oligos were immobilized onto FC-1, FC-2, FC-3 and FC-4 with RUs 94, 97, 92 and 98, respectively. The buffer used for all the experiments was 10 mM Tris-HCl, pH 7.4, 100 mM NaCl, 1mM EDTA containing 0.1% Tween 20 and 0.03% glycerol. To measure the binding, 100 or 200 nM of Rv0081-wt protein and 100-3200 nM for Ala (Rv0081 S/Q-A), and 100-6400 nM for Asp mutants (Rv0081 S-D) were flown over the chip. Binding assays were performed using flow rates of 50 μl/min; association was measured for 120 sec, while dissociation was measured for 180 sec. After each cycle, the chip was regenerated with consecutive injections of 1M NaCl. The SPR data was evaluated using the BIAevaluation software version 3.2 (Biacore). Three sets of data were collected using two different SA chips.

#### Kinetics Study

Kinetic measurements were performed by flowing various concentrations of Rv0081-Wt and Rv0081-S/Q-A proteins over the SA-chip immobilized with the Wt and M2-biotinylated oligos. The data collected were fitted with suitable kinetic models, i.e. either with Bi-valent model for wild-type oligo sequences or 1:1 Langmuir model for the mutant oligo sequences using the BIAevaluation software version 3.2 (Biacore).

### M. tuberculosis growth and stress conditions

Mtb H37Rv strain was used in the present study. Mycobacterial growth was achieved as described earlier(45). Mtb was grown in 7H9 media (Himedia, India) supplemented with 10% OADC media (Himedia, India), 0.4 % glycerol, 0.05 % Tween-80, at 37°C until its OD_600_ reached 0.6 to 0.8. Mtb cultures were then given hypoxic stress as described earlier described earlier with few modifications (23, 45). In brief, hypoxic stress was given by (i) low aeration (without shaking) or (ii) low aeration by covering with a layer of mineral oil (without shaking) or (iii) treatment of culture with 250 μM nitric oxide (NO). The cultures were harvested by centrifugation at 2465x g for 5 min in 4°C both pre and post stress. Pellet was resuspended in wash buffer [50 mM Tris-HCl pH 7.4, 150 mM NaCl, 1 mM EDTA, 5% glycerol and 1x protease and phosphatase inhibitor mix (Roche)] and centrifuged at 4°C at 2465x g and the pellet was stored at −80°C after flash-freezing in liquid-nitrogen. Each of the culture samples were checked for contamination before and after stress by using Ziehl–Neelsen staining (Himedia, India). The experiment in each condition was carried out in duplicate.

### Co-Immunoprecipitation (Co-IPs) of Rv0081 with its interacting partners

The cell pellet of 35 ml Mtb culture grown in normal or stress conditions were resuspended in 1 ml lysis buffer (50 mM Tris-HCl pH 7.4, 150 mM NaCl, 1 mM EDTA, 1% NP-40, 5% glycerol) and 1x protease and phosphatase inhibitor (Roche). Cells were lysed by bead beating with 0.1 mm glass beads for 10 cycles of 1 min on and 1 min off cycle. Cell lysates were clarified by centrifugation at 13800 x g for 20 min and filtered through 0.22 µm syringe filter (Merck). The total protein amounts in cell lysates were estimated by Bradford reagent (Biorad). About 180 µg of cell lysates from four different conditions were precleared by mixing about 20 µl of protein A/G beads (Pierce) and incubation for 4-5 hrs with end to end rotation of tubes. Lysates were centrifuged at 1500 rpm for 5 min and supernatants were transferred to new tubes. Lysates were mixed with ~30 µl protein A/G beads bound with approx. 5 μg of polyclonal Rv0081 antibody (Screening of poly-Ab Rv0081 was done by western blot using purified Rv0081-wt and further validated by pulling down native Rv0081 from cell lysate using mass-spectrometry) or 1 µg of IgG as a control (data not shown) and incubated at 4 °C for overnight with end to end rotation of tubes. Bound beads were washed 3 times with 200 µl of ice-cold lysis buffer. About 30 μl of 2x Laemmli SDS-PAGE loading buffer was added to each prep and were heated at 95 °C for 5 min. Tubes were centrifuged briefly and separated on 12.5 % SDS-PAGE. Proteins were transferred to PVDF membrane for Western blot analysis. The blotted membranes were blocked with blocking buffer [3% BSA in 1x Tris-buffered saline (TBS) with 0.1% Tween-20] for 2 hr. About 2 µg of phosphoserine mAb was added in 4 ml of blocking buffer and incubated overnight at 4 °C. Blots were probed with HRP-conjugated anti-mouse IgG (1:3000 dilutions). Signals were developed using Horseradish peroxidase (HRP)-chemiluminescence kit (Invitrogen) and imaged using BioRad imager. This experiment was repeated twice.

All experiments involving mycobacteria were performed at SB laboratory approved for mycobacterial work (UH/SLS/IBSC/SB/BSL-F-60) by University of Hyderabad Institutional Biosafety Committee constituted by Department of Biotechnology, Govt. of India.

## ACKNOWLEDGEMENTS

The funding was provided by Department of Biotechnology, Ministry of Science and Technology grants BT/PR3260/BRB/10/967/2011 and BT/PR15450/COE/34/46/2016.

We thank Dr. Thomas C. Zahrt, DMMG, Center for Infectious Disease Research, Medical College of Wisconsin USA for providing Rv0081 construct plasmid. We thank Dr. Saikrishanan (IISER Pune XRD, facility) and beamline staff at the Elettra synchrotron Trieste (XRD1) for their assistance during data collection. We thank Ms. Tanuja N Bankar, Dr. Krishnasastry and Dr. Ramanmurthy for generation of poly-Ab of Rv0081 in NCCS animal house facility, Dr. C.M Santosh Kumar for his suggestion in Rv0081 purification and Ms. Kriti Chopra to help in preparing phylogenetic trees. The authors acknowledge financial assistance from Indian Council of Medical Research, New Delhi (AK), Council of Scientific Industrial Research, New Delhi (SP and HSP).

## ABBREVATIONS

Mtb: *Mycobacterium tuberculosis*
SPR: surface Plasmon resonance
MSA: multiple sequence alignment
TB: tuberculosis
RNR: ribonucleoside reductases
FHL: formate hydrogen lyase

